# Variability of in vivo sarcomere length measures in the upper limb obtained with second harmonic generation microendoscopy

**DOI:** 10.1101/2021.11.17.468984

**Authors:** Amy N Adkins, Ryan Fong, Julius P A Dewald, Wendy M Murray

## Abstract

The lengths of a muscle’s sarcomeres are a primary determinant of its ability to contract and produce force. In addition, sarcomere length is a critical parameter that is required to make meaningful comparisons of both the force-generating and excursion capacities of different muscles. Until recently, in vivo sarcomere length data have been limited to invasive or intraoperative measurement techniques. With the advent of second harmonic generation microendosopy, minimally invasive measures of sarcomere length can be made for the first time. This imaging technique expands our ability to study muscle adaptation due to changes in stimulus, use, or disease. However, due to the inability to measure sarcomeres outside of surgery or biopsy, little is known about the natural, anatomical variability in sarcomere length in living human subjects. To develop robust experimental protocols that ensure data provide accurate representations of a muscle’s sarcomere lengths, we sought to quantify experimental uncertainty associated with in vivo measures of sarcomere lengths. Specifically, we assessed the variability in sarcomere length measured 1) within a single image, along a muscle fiber, 2) across images captured within a single trial, across trials, and across days, as well as 3) across locations in the muscle using second harmonic generation in two upper limb muscles with different muscle architectures, functions, and sizes. Across all of our measures of variability we estimate that the magnitude of the uncertainty in in vivo sarcomere length are on the order of ~0.25μm. In the two upper limb muscles studied we found larger variability in sarcomere length within a single insertion than across locations. We also developed custom code to make measures of sarcomere length variability across a single fiber and determined that this codes’ accuracy is an order of magnitude smaller than our measurement uncertainty due to sarcomere variability. Together, our findings provide guidance for the development of robust experimental design and analysis of in vivo sarcomere lengths in the upper limb.

## Introduction

Whole muscle is made up of hundreds of thousands of sarcomeres arranged in series and parallel. The length of a muscle’s sarcomeres is a primary determinant of a muscle’s ability to contract and produce force (Gordon, Huxley et al. 1966). In addition, when quantifying muscle architecture, sarcomere length is a critical parameter that is required to make meaningful comparisons of both the force-generating and excursion capacities of different muscles. With the advent of second harmonic generation (SHG) microendoscopy for measuring sarcomere length minimally invasively (Llewellyn, Barretto et al. 2008), there are exciting and novel opportunities to measure sarcomere length *in vivo*.

Anatomical studies in cadavers, which involve dissection and measurement across multiple scales (whole muscles, fascicles, and sarcomeres), demonstrate that a major distinction among different skeletal muscles in the human body is the number of sarcomeres in series and parallel (Brand, Beach et al. 1981, Lieber, Fazeli et al. 1990, Lieber and Fridén 2000). Serial sarcomere number (SSN) is a particularly important architectural parameter, as it describes the range of lengths over which a muscle can actively generate force and, given optimal sarcomere length, a measure of the fascicle length at which the muscle will produce its maximum isometric force. The number of parallel sarcomeres is proportional to the maximum isometric force a muscle can produce. In anatomical studies, SSN is commonly characterized by a muscle’s optimal fascicle length (OFL), which is calculated as the ratio of fascicle and sarcomere lengths measured from a muscle’s dissected fascicles, multiplied by optimal sarcomere length. Similarly, the number of sarcomeres in parallel is characterized by a muscle’s physiological cross-sectional area (PCSA), which is calculated from the ratio of the muscle’s volume and OFL, with a correction for the pennation angle of the fibers to estimate how much of the force-generating capacity is transmitted by the tendon. Measures of *in vivo* fascicle length and muscle volume have been made using safe and effective imaging techniques for decades. The novel capacity to make minimally invasive measures of sarcomere length *in vivo,* provides new opportunities to study functionally meaningful muscle parameters (OFL and PCSA) and how they vary in living subjects across muscles, individuals, and due to alterations in muscle stimulus or use.

Due to the novelty of methods for sarcomere length measurement that do not involve surgery or biopsy, little is known about the natural, anatomical variability in sarcomere length in living human subjects. Such an understanding is critical for the development of robust experimental protocols that ensure data provide accurate representations of a muscle’s sarcomere lengths. Animal muscle studies (e.g.(Moo, Fortuna et al. 2016, Moo, Leonard et al. 2017)) and a single *in vivo* study on the tibialis anterior (Lichtwark, Farris et al. 2018) suggest that sarcomere length can vary considerably throughout a muscle. Importantly, the SHG microendoscopy method provides only a small field-of-view (82μm × 82μm, ~20-35 sarcomeres in series) (Llewellyn, Barretto et al. 2008, Sanchez, Sinha et al. 2015). In addition, current protocols for processing SHG data utilize a method which provides a single measure - the mean sarcomere length - per image, precluding information about within fibre variability. To enable effective studies of how muscle sarcomere length may differ or change (e.g., across limbs, after an exercise intervention, following an injury, etc.), more information describing the natural anatomical variability, both across different locations in a single muscle and along a single muscle fiber, is needed.

In this study we aim to quantify experimental uncertainty associated with *in vivo* measures of sarcomere lengths made using second harmonic generation microendoscopy in two muscles of the upper limb (biceps brachii and flexor carpi ulnaris). Given the precision of the SHG imaging and measurements procedures established in the literature is small (~30 nm) (Sanchez, Sinha et al. 2015), we expect the primary sources of measurement uncertainty to include both natural anatomical variability of the lengths of the sarcomeres within these muscles and the variability associated with our test-retest reliability using this method. In this study, we assessed the variability in sarcomere length measured 1) within a single image, along a muscle fiber, 2) across images captured within a single trial, across trials, and across days, as well as 3) across locations in the muscle. Quantifying the degree of experimental uncertainty for sarcomere length measures made using second harmonic generation microendoscopy will aid interpretation of these novel data, guide the design of future work aimed at detecting sarcomere length differences among different populations, and enable error propagation when sarcomere length measures are incorporated with other anatomical measures to calculate functionally meaningful muscle architectural parameter values.

## Methods

### Data collection

The 14 participants enrolled in this study provided informed consent; Northwestern University’s Institutional Review Board approved this study’s procedures. To image sarcomeres *in vivo*, a microendoscopic needle was inserted either into the long head of the biceps brachii or the flexor carpi ulnaris with its optical lenses aligned parallel to the fascicle direction. Insertion of the probe was guided through palpation techniques and ultrasound imaging. A microscope (Zebra Medical Technologies, Palo Alto, CA) which uses second-harmonic generation to capture the intrinsic striation pattern (A-bands) of sarcomeres was attached to the needle for imaging. Images were collected at 1.9Hz for about 2-5minutes (~250-600 images).

Imaging for the both the biceps brachii and FCU was done under passive conditions with participants seated. For measures on the biceps the participants arm was placed in shoulder abducted 85°, elbow extended 25°, and wrist at neutral (0°) which was verified by handheld goniometric measures. Similarly, imaging of the FCU was with the elbow at 70° of flexion and the wrist at neutral (0°).

#### 1 Variability along fiber

Current image processing methods for analysis of SHG images (mean sarcomere length code; MSLC) report a single mean sarcomere length value for an entire image (e.g., (Sanchez, Sinha et al. 2015, Adkins, Dewald et al. 2021). Here, we developed and validated an image processing and analysis code (individual sarcomere length code: ISLC) which quantifies: the lengths of individual sarcomeres within an image, the mean length of these sarcomeres, and standard deviation in sarcomere length within the image. To verify the performance of the ISLC, realistic virtual phantoms were designed, tested for their similarity to real data, and then used to calculate and compare the accuracy and precision of the novel ISLC and the existing MSLC.

### Individual Sarcomere Length Code

The image processing protocol developed in this study was designed to quantify the length of individual sarcomeres within an image, along the imaged fiber direction, using custom written MATLAB code. Specifically, the ISLC takes a Fast Fourier Transform (FFT) of each image and applies a threshold to reduce noise and clear the edges of the Fourier space. The inverse FFT (IFFT) is computed, and Canny edge detection is utilized to select the largest area in the image with sarcomeres (Figure 1A). Within the selected region, the fiber orientation is determined by binarizing the region and calculating the orientation relative to the horizontal axis of the box (Figure 1B). Sarcomere length is calculated along a single column of pixels as the pixel distance between pixel intensity peaks (A-bands), converted to microns (0.16μm per pixel) and multiplied by the cosine of the detected angle to correct for angle of fiber orientation (Figure 1C). This analysis process is repeated for distinct columns of pixels at 2μm intervals to enable the collection of sarcomere length data from separate myofibrils (Lieber 2002, Moo, Fortuna et al. 2016).

**Figure 1:**
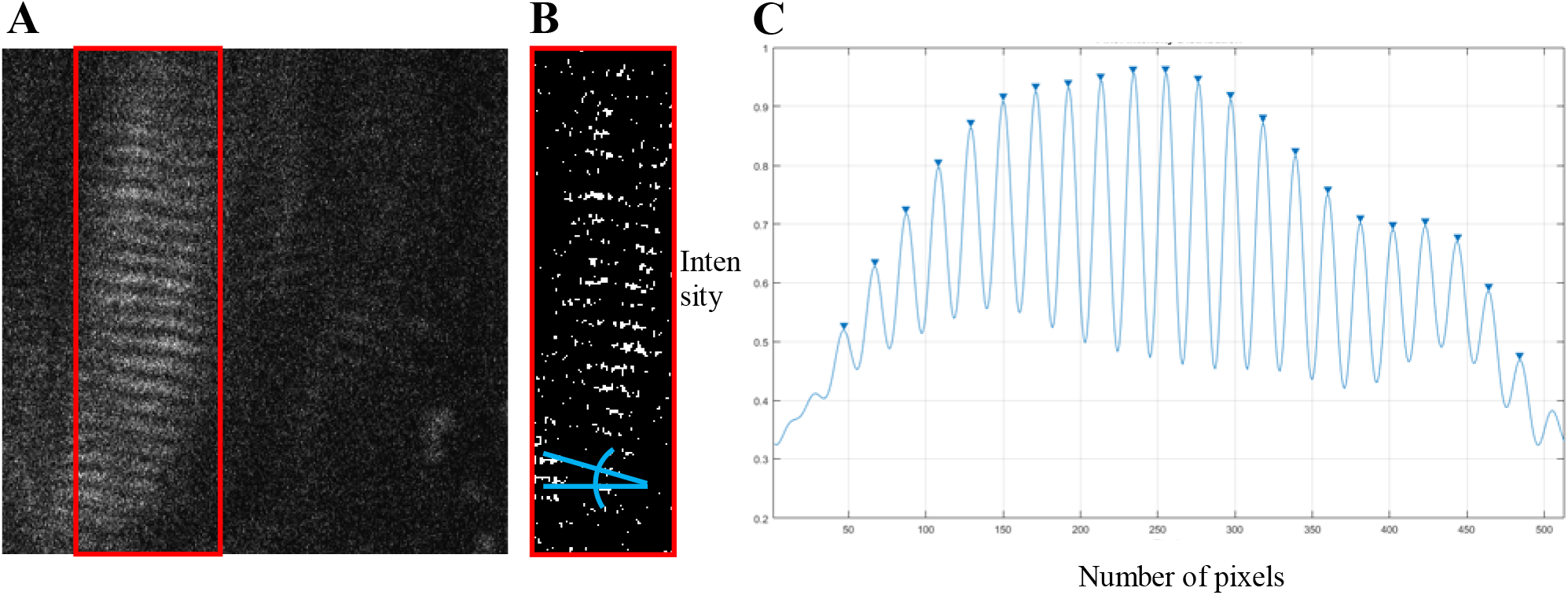
(A) Sample raw SHG image of *in vivo* biceps brachii sarcomere data (512 × 512 pixels, 82 × 82 μm). Canny edge detection selected region enclosed by red box. (B) Binarized selected region with blue lines indicating the angle relative to horizontal. (C) Pixel versus intensity distribution where A-bands are represented by peaks

### Realistic virtual Phantoms

Virtual phantoms were designed to closely mimic raw, *in vivo* data. A range of phantoms with sarcomere length averages and standard deviations representative of cadaveric data from the biceps brachii (data from (Murray, Buchanan et al. 2000)) were generated using a series of normally distributed sinusoids (Figure 2A). To mimic observed raw SHG data, the image signal in the phantom was degraded by removing 70% of the pixels at random. A sinusoidal geometric transformation was applied to create fiber bending, and Gaussian white noise was added to capture similar signal quality (Figure 2B-C). To assess the phantoms’ similarity to real data, the signal-to-noise ratio (SNR) for the phantoms was measured and then adapted to mimic the range of SNR values found in raw SHG images from the biceps brachii.

**Figure 2:**
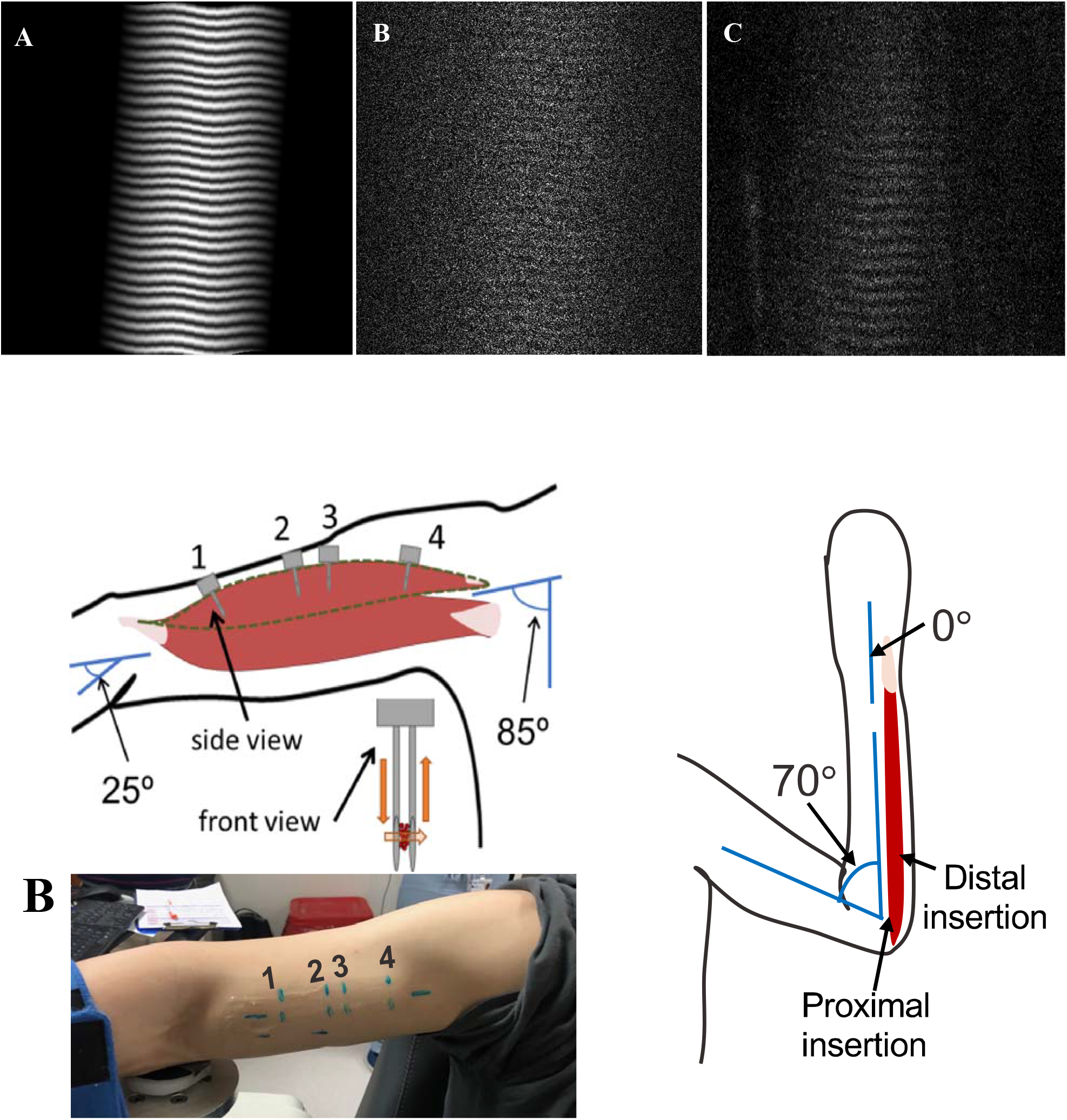
(A) Virtual Phantom created with real sarcomere lengths and variability, distortion, and rotation. (B) Realistic Virtual Phantom with added noise. (C) Representative raw data from *in vivo* biceps brachii.

### Determining Accuracy and comparing ISLC and MSLC

Using 300 virtual phantom images with known sarcomere lengths, and varying image quality, the code’s accuracy and precision were determined by comparing the ISLC output to the known phantom values. The accuracy and precision of the ISLC code developed in this study was defined from the differences between the “true” mean sarcomere lengths defined in the virtual phantoms and the mean sarcomere length values output from the ISLC. Accuracy was defined by the difference in means; precision was defined by the standard deviation of the differences. This analysis was repeated for the MSLC to enable direct comparison of the two methods. A Bland-Altman test was implemented to determine agreement between sarcomere lengths obtained using ISLC and sarcomere measurements obtained using MSCL.

#### 2 Variability across images, trials, and days

Sarcomere length variability across different images from a single imaging trial (defined as a single needle insertion and microscope attachment) was assessed using the standard deviation of the mean sarcomere length from all processed images in the trial. For this study, we analyzed images that were acquired in the non-paretic biceps of 7 individuals with chronic hemiparetic stroke and in the biceps from both the dominant and non-dominant limbs of 4 individuals with no neuromuscular impairments (Table 4.1). Therefore, SHG images were collected from the long head of the biceps brachii in 15 limbs. On average, we collected 145 ± 108 images per trial. Mean sarcomere length per image was calculated using MSLC.

**Table 4.1.**
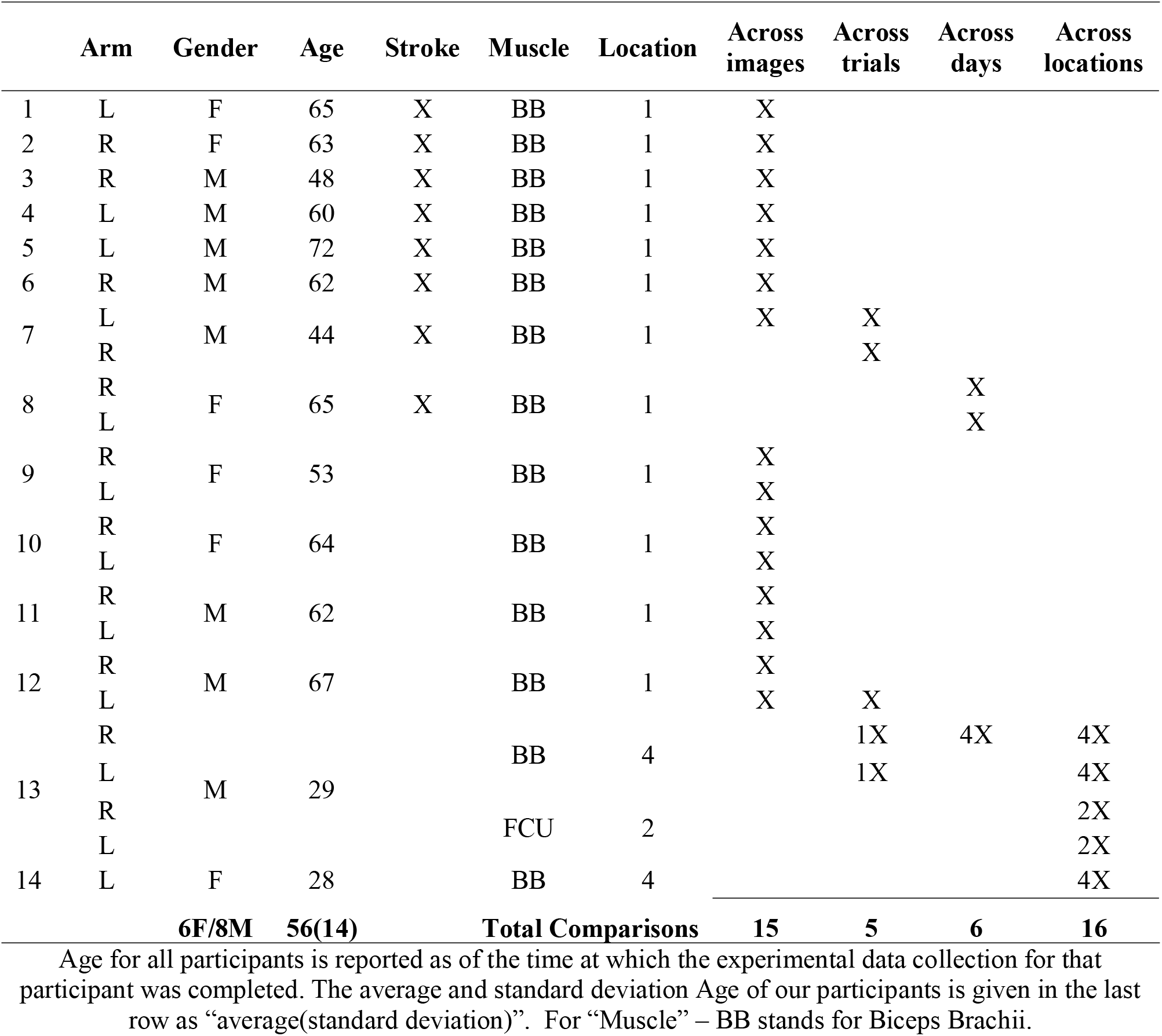
Participant demographic and participation information

Images collected from 3 participants were analyzed to evaluate the variability in sarcomere lengths measured when two trials of data were collected in the same muscle (Table 4.1, different trials arise from a single needle insertion, but distinct microscope attachment). Across trials, there is slight movement of the probe within the muscle due to the removal and reattachment of the microscope on the probe. Because the needle is not removed and reinserted, we expect we are likely sampling from the same relatively small, localized area, but from different sarcomeres due to the probe movement. In total, we analyzed two trials of data collected from 5 distinct insertions, including insertions in both arms of two participants, and a single arm from a third participant. Variability between 2 trials of image data was quantified as the difference in the average sarcomere length per trial (i.e., the average value of the output of MSLC for all the images in that trial)

The variability in sarcomere lengths measured on different days (a minimum of 1 week apart) was assessed in the long head of the biceps brachii. Images collected from a single needle insertion in both limbs of 1 participant and from 4 insertion sites (at different locations in the muscle) from a single limb in a second participant were analyzed (Table 4.1). Because measurements were taken on different days the insertion locations were not in the exact location on Day 2 as on Day 1, but the investigator’s intention was to insert the needle consistently across days. Data was collected a minimum of 1 week apart. Variability in the sarcomere lengths measured on 2 different days was quantified as difference in the average sarcomere length captured on “Day 1” and “Day 2” (i.e., the average value of the output of MSLC for all the images in the single trial of data collected from a given needle insertion location each day). In four of the repeated measures across days, the insertion location was marked on the limb and the distance between insertion locations was measured and determined to be less than 1 cm apart (average ± standard deviation, 0.54 ± 0.17 cm).

#### 3 Variability in different locations in the muscle

Sarcomere lengths in the biceps brachii were measured at distinct locations in the muscle in three arms of two individuals (1 female, 28yrs; 1 male, 26yrs) (Figure 3). Four insertions were made; insertions near the distal and proximal ends of the muscle belly (position 1 and 4), and two insertions in the center of the muscle belly approximately 1cm apart (positions 2 and 3). In both arms of a single individual (male, 29 years old) sarcomere length measures were made at two locations along the flexor carpi ulnaris muscle; a proximal location where the fibers run longitudinally in the muscle, and a distal location where the fibers are pennated (Figure 3). Measurements of fascicle length of the long head of the biceps brachii and the flexor carpi ulnaris were obtained using extended field-of-view ultrasound (Acuson S2000 Ultrasound System, Siemens Medical Solutions USA, Inc., Mountain View, CA) at the same joint posture sarcomere length was obtained (Figure 4). Ultrasound images were exported as uncompressed DICOM files and fascicles were measured using the segmented line tool in ImageJ (Wayne Rasband, National Institutes of Health, Bethesda,MD).

**Figure 3.** A) schematic of arm position and location of probe insertion in the long head of the Biceps Brachii. Illustration of probe showing the direction of laser light and interaction with muscle tissue (orange). B) Picture of location of four probe insertions in one of the study participants. C) Schematic of the elbow and wrist position and locations of the probe insertion in the flexor carpi ulnaris.

**Figure 4.**
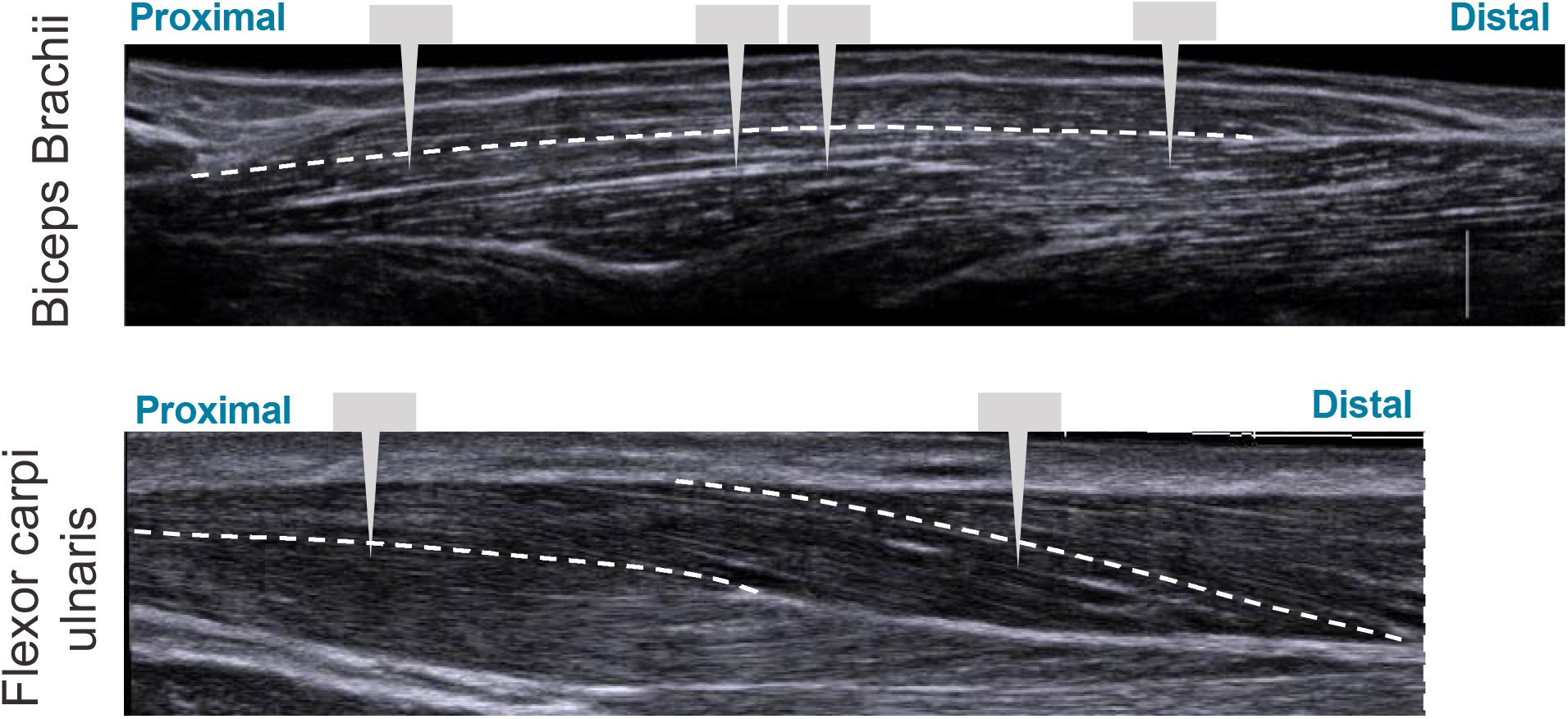
Illustration showing extended field-of-view ultrasound images with approximate locations of probe insertions along the muscle. White dashed line represents fascicle paths.

## Results

### 1 Variability along fiber

The ISLC code developed in this study was accurate to 0.02μm and precise to 0.02μm. While the analysis from the virtual phantoms indicates the MSLC code was slightly less accurate than the ISLC (accuracy 0.03μm, precision 0.02μm), the Bland-Altman analysis (Figure 5) demonstrates that mean sarcomere lengths that result from the two analyses are comparable. There is a minimal bias between the two methods (−0.0025μm; 95% limits-of-agreement = −0.0809μm - 0.0759μm), there is no observable systematic variance, and the measurements lay within the limits-of-agreement.

**Figure 5.**
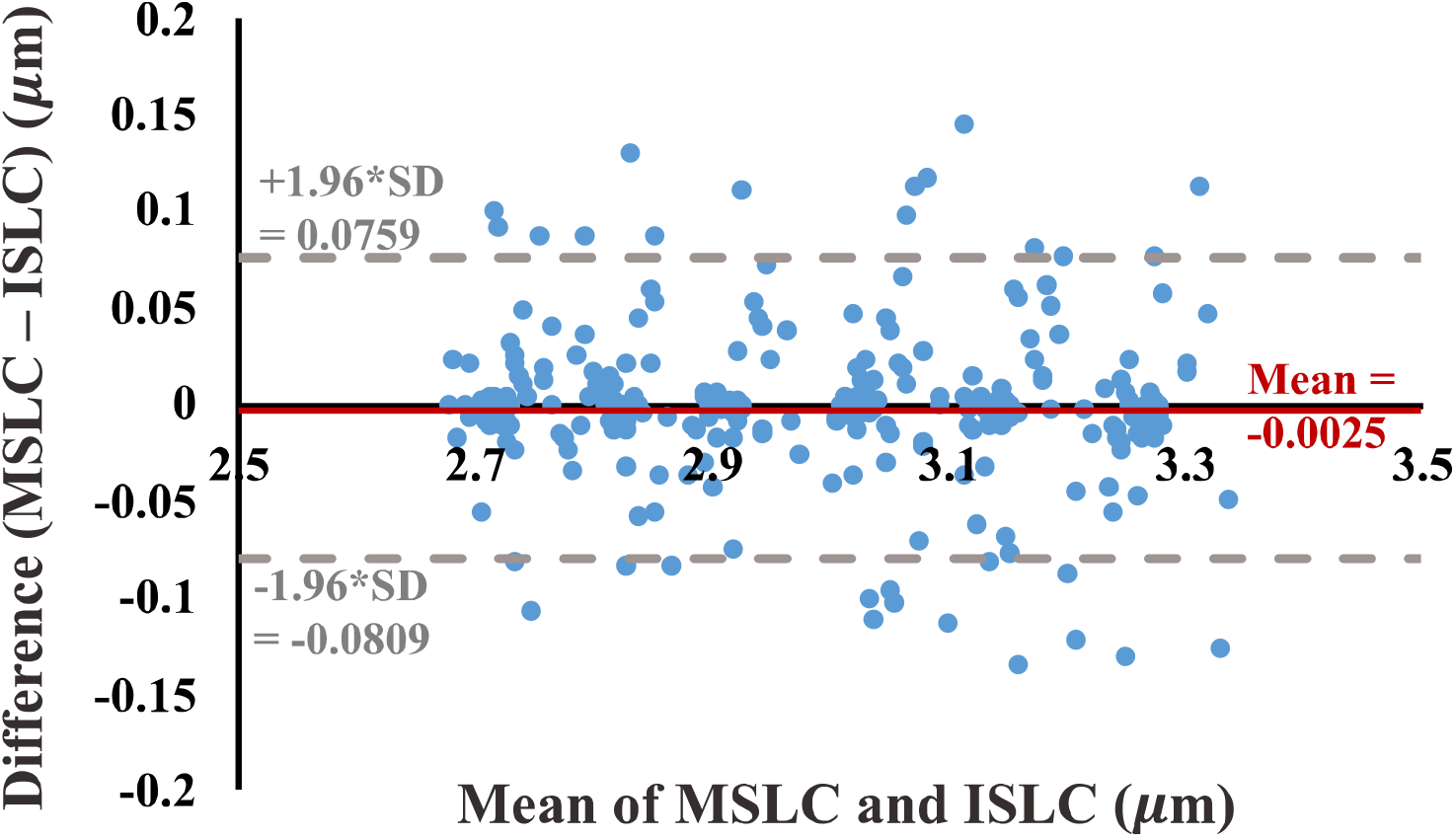
Bland-Altman test of agreement between our individual sarcomere length code (ISLC) and the image processing method currently utilized in the literature, MSLC. The x-axis is the average of the MSLC and ISLC sarcomere length measurements and the y-axis is the difference between the sarcomere lengths calculated from the two methods. The solid red line indicates the bias (−0.0025μm) and the dashed grey lines represent the lower (−0.0809μm) and upper (0.0759μm) limits-of-agreement (mean difference ± 1.96*standard deviation of the difference).

### 2 Variability across images, trials, and days

The mean sarcomere length observed in a single image varied across the multiple images that are captured in a single imaging trial (ie, single needle insertion and microscope attachment), although this variability is relatively small. Across 15 trials, the standard deviation of the mean sarcomere length per image within a trial was 0.20 microns ± 0.07 microns. The magnitude of the difference in mean sarcomere length measured in different trials (average ± standard deviation: trials: 0.18 ± 0.10μm) was comparable to the variability across images and the difference in mean sarcomere length measured in the same muscle on different days (0.15 ± 0.13μm).

### 3 Variability across muscle

The uncertainty in sarcomere length that was introduced by sampling from various, distinct muscle locations was not larger than the variability in mean sarcomere length we observed across images within a single trial (i.e. needle insertion). For the biceps brachii, the difference in median sarcomere length across four locations was 0.15 ± 0.12μm. For the flexor carpi ulnaris, the difference in median sarcomere length across two locations was 0.13 ± 0.09μm. Standard deviations of images collected in a single trial and location averaged 0.18 ± 0.07μm for the biceps and 0.36 ± 0.19μm for the FCU. The largest difference in median sarcomere length between any two locations in a single muscle occurred in the biceps brachii of arm 3 between the proximal and center-proximal location (difference = 0.43μm). Notably, for both muscles and for all arms evaluated, the maximum difference in median sarcomere length observed at different locations (c.f. Figure 6 & 4.7 purple shaded region) was less than the range of sarcomere lengths observed in a single stick (colored probability distribution of each violin plot for each arm).

**Figure 6.**
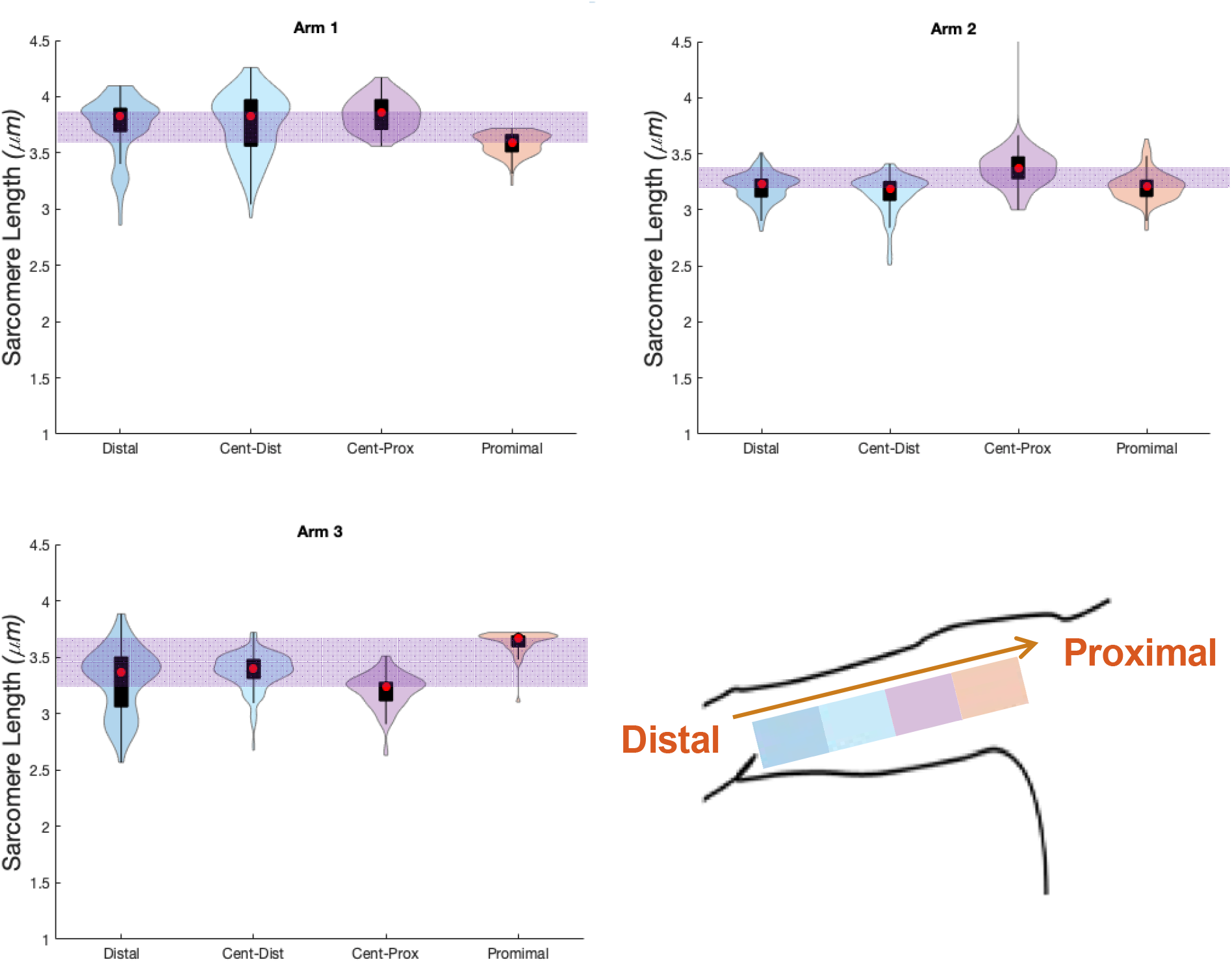
Violin plots for 4 locations, and three arms showing median of data (red circle), interquartile range (black box plot), and probability density of data at each location (shaded violin shape).

**Figure 7.** Schematic of the two insertion locations (left) in the flexor carpi ulnaris., and three arms showing median of data (red circle), interquartile range (black box plot), and probability density of data at each location (shaded violin shape).

## Summary/ functional implications

Based on the sources of variability in sarcomere length measures in the upper limb muscles investigated in this study, we estimate the uncertainty in sarcomere length associated with collecting SHG microendoscope images in a single imaging session from a single needle insertion within the muscle to be on the order of 0.25μm (Figure 8), which was ~7-8% of the average sarcomere lengths in the biceps brachii and FCU. For two muscles in the upper limb that have different functions, size, and architectures, the variability in mean sarcomere length observed across different images and collected from a single needle insertion was of a comparable magnitude to the variability of measures observed on different days or at different locations in the muscle. Notably, this level of uncertainty (~0.25μm) is an order of magnitude greater than the improvement in measurement accuracy obtained if the variability in sarcomere length within an image is considered along a muscle fiber (difference in accuracy between ISLC and MCLC ~0.01μm). Given it is used in the calculation of functionally relevant architectural parameters, the uncertainty in sarcomere length measures we observed would yield uncertainty in calculations of optimal fascicle length and physiological cross-sectional area of ~6.7 – 7.7%, independent of any experimental uncertainty in measurements of fascicle length or muscle volume.

**Figure 8.**
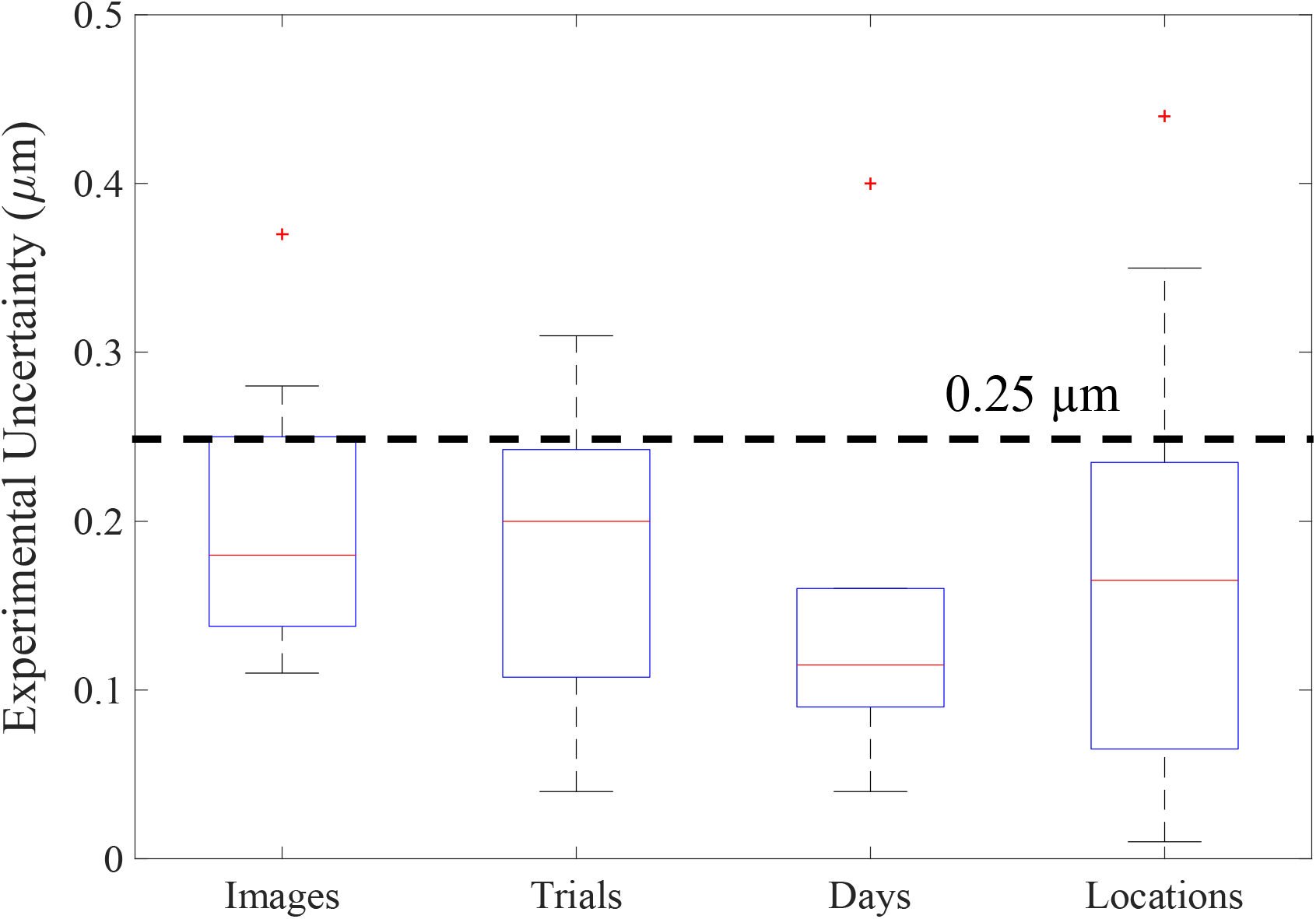
Boxplots of the various sources of experimental uncertainty investigated in this study. The red line represents the median, the blue box extends from the 25^th^ to the 75^th^ percentile of the data, and the dashed error bars, referred to as whiskers, represent the range of all data excluding outliers (red “+”). For images the y-axis represents the standard deviation in mean sarcomere length across images. For all other x-axis elements, the y-axis represents the absolute difference in mean sarcomere length between trials, days, and locations.

## Discussion

We investigated both anatomical and experimental sources of variability in sarcomere length measures obtained in vivo, in the upper limb, using second harmonic generation microendoscopy. Based on assessments completed in two muscles in the upper limb that have different functions, size, and architectures, we estimate the magnitude of the uncertainty in such measures to be on the order of ~0.25μm. Notably, we found that, in both a fusiform muscle (biceps brachii) and pennate muscle (flexor carpi ulnaris) of the upper limb, variability in sarcomere length within a single insertion is larger than the variability across locations in these muscles. We developed custom image processing code which takes measures of individual sarcomeres in an image, allowing for the quantification of sarcomere length variability along a muscle fiber and across myofibrils in a localized area (82μm by 82μm). We found that the uncertainty in mean sarcomere length measures made using this ISLC was comparable to current processing methods (MSLC) and an order of magnitude smaller (~0.02μm) than the uncertainty in sarcomere length of all other parameters investigated in this study (i.e., across images, trials, days, locations). Together, our findings provide guidance for the development of robust experimental design and analysis of in vivo sarcomere lengths in the upper limb.

The uncertainty in sarcomere length - and therefore the most functionally meaningful muscle parameters (optimal fascicle length and PCSA) - found in this study (<8%) is smaller than differences found in *in vivo* sarcomere length and fascicle length of various studies in the literature; suggesting that using SHG to detect *in vivo* sarcomere length adaptation is practical. For example, intraoperative measures of sarcomeres have been shown to be much longer in children with cerebral palsy (an injury of the brain during development) than sarcomere length estimates for (Mathewson, Ward et al. 2015) (~88%) or intraoperative measures (Lieber and Fridén 2019) (~46%) in typically developing children. In addition, following three weeks of Nordic hamstring training, a 17% increase in sarcomere length was found in the biceps femoris long head (Pincheira, Boswell et al. 2021). Due to the relative ease of obtaining fascicle length measures as compared to sarcomere length measures in vivo, many studies have reported adaptation of fascicle length to altered use or stimulus. Previous studies in elbow flexors of individuals with chronic hemiparetic stroke report substantial decreases in fascicle length in extended joint postures (~19% decrease in biceps brachii at 25º elbow flexion (Nelson, Murray et al. 2018), 15% decrease in brachialis at 10º elbow flexion (Li, Tong et al. 2007)). Fascicle length in gastrocnemius medialis in individuals who regularly wear high heels were found to be 11% shorter than non-high heel wearers (Csapo, Maganaris et al. 2010). If these changes in fascicle length were due solely to changes in sarcomere length (i.e., a decrease in sarcomere length of 15-19%), our study suggests SHG microendoscopy would be able to reliably detect such changes. Notably, in a single individual who underwent a leg lengthening surgery (4cm distraction) to correct a limb discrepancy, sarcomere length decreased by ~15% while fascicle length increased by over 100% (Boakes, Foran et al. 2007). Together, these changes resulted in an overall increase in serial sarcomere number of 134.6% (Boakes, Foran et al. 2007), emphasizing the importance of measurement of both fascicle and sarcomere length to fully understand muscle function and plasticity.

Our data suggest the approach of measuring sarcomere lengths from a single insertion site in the muscle belly is appropriate for an understanding of average sarcomere length and variability in a fusiform (biceps brachii) and pennate (flexor carpi ulnaris) muscles of the upper limb. This finding will guide the development of future studies determining interlimb or interindividual differences in sarcomere length in the biceps brachii or flexor carpi ulnaris. For example, an a priori power analysis to determine the difference in sarcomere length between two dependent means (e.g., left and right arm) using SHG microendoscopy (standard deviation of 0.25 micrometers) would demonstrate that a power over 0.8 would be achieved with 10 participants with a detectable difference of 0.25 micrometers (effect size of 1) between limbs.

This study provides insight which will be invaluable for the design, implementation, and analysis of future in vivo architectural studies aiming to explore muscle plasticity. However, there are several limitations in this work which should be considered. In particular, the relatively small cohort of individuals we evaluated limited our ability to make statistical conclusions. In addition, we only explore sarcomere length variability in a small number of muscles and at a single joint posture and contraction state per muscle. However, most previous in vivo muscle architectural studies demonstrate differences in fascicle length without normalizing to sarcomere length or make estimates of PCSA by using this non-normalized fascicle length. With the addition of sarcomere length differences across populations, following interventions, or impairment of the two most functionally relevant parameters of muscle (serial sarcomere length and PCSA) can be explicitly calculated. This work provides confidence in the ability to use SHG microendoscopy to make measures of in vivo sarcomere length to detect difference in functionally relevant muscle parameters on the order of 8% or greater.

## Figure Legends

## Author Contributions

A.N.A., J.P.A.D., and W.M.M. designed research; A.N.A., R.F, J.P.A.D, and W.M.M. performed research; A.N.A., R.F, and W.M.M. analyzed data; and A.N.A., R.F, J.P.A.D, and W.M.M. wrote the paper.

## Acknowledgments

We would like to thank the study participants, Vikram Darbhe and Jorie Budzikowski for assistance in data collection, and Zebra Medical Technologies (now Enspectra Health) for their support with data collection and image processing. This work is supported by the National Science Foundation (NSF) Graduate Research Fellowship Program under Grant No. DGE-1324585, as well as National Institutes of Health (NIH) R01 D084009 and F31 AR076920. Any opinions, findings, and conclusions or recommendations expressed in this material are those of the authors and do not necessarily reflect the views of the NSF or NIH.

## Data Availability Statement

The datasets generated for this study can be found in Arch, Northwestern University Research and Data Repository [LINK] upon publication.

## References

Adkins, A. N., J. P. A. Dewald, L. P. Garmirian, C. M. Nelson and W. M. Murray (2021). “Serial sarcomere number is substantially decreased within the paretic biceps brachii in individuals with chronic hemiparetic stroke.” Proceedings of the National Academy of Sciences 118(26): e2008597118.

Boakes, J. L., J. Foran, S. R. Ward and R. L. Lieber (2007). “CASE REPORT: Muscle Adaptation by Serial Sarcomere Addition 1 Year after Femoral Lengthening.” Clinical orthopaedics and related research 456: 250–253.

Brand, P. W., R. B. Beach and D. E. Thompson (1981). “Relative tension and potential excursion of muscles in the forearm and hand.” Journal of Hand Surgery 6(3).

Csapo, R., C. N. Maganaris, O. R. Seynnes and M. V. Narici (2010). “On muscle, tendon and high heels.” The Journal of Experimental Biology 213(15): 2582–2588.

Gordon, A., A. F. Huxley and F. Julian (1966). “The variation in isometric tension with sarcomere length in vertebrate muscle fibres.” The Journal of physiology 184(1): 170–192.

Li, L., K. Y. Tong and X. Hu (2007). “The Effect of Poststroke Impairments on Brachialis Muscle Architecture as Measured by Ultrasound.” Arch Phys Med Rehabil 88: 243–250.

Lichtwark, G. A., D. J. Farris, X. Chen, P. W. Hodges and S. L. Delp (2018). “Microendoscopy reveals positive correlation in multiscale length changes and variable sarcomere lengths across different regions of human muscle.” Journal of Applied Physiology 125(6): 1812–1820.

Lieber, R. L. (2002). Skeletal muscle structure, function & plasticity: The physiological basis of rehabilitation. Philadelphia, Lippincott Williams & Wilkins.

Lieber, R. L., B. M. Fazeli and M. J. Botte (1990). “Architecture of Selected Wrist Flexor and Extensor Muscles.” Journal of Hand Surgery-American 15A(2): 244–250.

Lieber, R. L. and J. Fridén (2000). “Functional and clinical significance of skeletal muscle architecture.” Muscle & Nerve: Official Journal of the American Association of Electrodiagnostic Medicine 23(11): 1647–1666.

Lieber, R. L. and J. Fridén (2019). “Muscle contracture and passive mechanics in cerebral palsy.” Journal of Applied Physiology 126(5): 1492–1501.

Llewellyn, M. E., R. P. J. Barretto, S. L. Delp and M. J. Schnitzer (2008). “Minimally invasive high-speed imaging of sarcomere contractile dynamics in mice and humans.” Nature 454(7205): 784–788.

Mathewson, M. A., S. R. Ward, H. G. Chambers and R. L. Lieber (2015). “High resolution muscle measurements provide insights into equinus contractures in patients with cerebral palsy.” Journal of Orthopaedic Research 33(1): 33–39.

Moo, E. K., R. Fortuna, S. C. Sibole, Z. Abusara and W. Herzog (2016). “In vivo Sarcomere Lengths and Sarcomere Elongations Are Not Uniform across an Intact Muscle.” Frontiers in Physiology 7(187).

Moo, E. K., T. R. Leonard and W. Herzog (2017). “In Vivo Sarcomere Lengths Become More Non-uniform upon Activation in Intact Whole Muscle.” Frontiers in Physiology 8(1015).

Murray, W. M., T. S. Buchanan and S. L. Delp (2000). “The isometric functional capacity of muscles that cross the elbow.” J Biomech 33(8): 943–952.

Nelson, C. M., W. M. Murray and J. P. A. Dewald (2018). “Motor Impairment–Related Alterations in Biceps and Triceps Brachii Fascicle Lengths in Chronic Hemiparetic Stroke.” Neurorehabilitation and Neural Repair 32(9): 799–809.

Pincheira, P. A., M. A. Boswell, M. V. Franchi, S. L. Delp and G. A. Lichtwark (2021). “Biceps femoris long head sarcomere and fascicle length adaptations after three weeks of eccentric exercise training.” bioRxiv: 2021.2001.2018.427202.

Sanchez, Gabriel N., S. Sinha, H. Liske, X. Chen, V. Nguyen, Scott L. Delp and Mark J. Schnitzer (2015). “In Vivo Imaging of Human Sarcomere Twitch Dynamics in Individual Motor Units.” Neuron 88(6): 1109–1120.

